# Identification of distinct genotypes in circulating RSV A strains based on variants on the virus replication-associated genes

**DOI:** 10.1101/2024.04.22.590570

**Authors:** Abdulafiz O. Musa, Sydney R. Faber, Kaitlyn Forrest, Kenneth P. Smith, Shaon Sengupta, Carolina B. López

## Abstract

Respiratory syncytial virus is a common cause of respiratory infection that often leads to hospitalization of infected younger children and older adults. RSV is classified into two strains, A and B, each with several subgroups or genotypes. One issue with the definition of these subgroups is the lack of a unified method of identification or genotyping. We propose that genotyping strategies based on the genes coding for replication-associated proteins could provide critical information on the replication capacity of the distinct subgroup, while clearly distinguishing genotypes. Here, we analyzed the virus replication-associated genes N, P, M2, and L from *de novo* assembled RSV A sequences obtained from 31 newly sequenced samples from hospitalized patients in Philadelphia and 78 additional publicly available sequences from different geographic locations within the US. In-depth analysis and annotation of the protein variants in L and the other replication-associated proteins N, P, M2-1, and M2-2 identified the polymerase protein L as a robust target for genotyping RSV subgroups. Importantly, our analysis revealed non-synonymous variations in L that were consistently accompanied by conserved changes in its co-factor P or the M2-2 protein, suggesting associations and interactions between specific domains of these proteins. These results highlight L as an alternative to other RSV genotyping targets and demonstrate the value of in-depth analyses and annotations of RSV sequences as it can serve as a foundation for subsequent *in vitro* and clinical studies on the efficiency of the polymerase and fitness of different virus isolates.

**IMPORTANCE:** Given the historical heterogeneity of Respiratory Syncytial Virus (RSV) and the disease it causes, there is a need to understand the circulating RSV strains each season. This requires an informative and consensus method of genotype definition. Here, we carried out a variant analysis that shows a pattern of specific variations among the replication-associated genes of RSV A across different seasons. Interestingly, these variation patterns point to previously defined interactions of domains within these genes, suggesting co-variation in the replication-associated genes. Our results also suggest a genotyping strategy that can prove to be particularly important in understanding the genotype-phenotype correlation in the era of RSV vaccination, where selective pressure on the virus to evolve is anticipated. More importantly, the categorization of RSV strains based on these patterns may prove to be of prognostic value.

## INTRODUCTION

Respiratory Syncytial Virus is a negative-sense single-stranded RNA virus that belongs to the family *Pneumoviridae* and order Mononegavirales. RSV circulates seasonally around the world, and it is estimated to infect every child once before the age of three, with the possibility of reinfections throughout life(1, 2). RSV leads to a wide variety of clinical outcomes that range from a mild cold to bronchiolitis, pneumonia, and death(2). RSV is a leading cause of hospitalizations in children and a major cause of morbidity in adults(3, 4). Each year, in the United States (US) alone, it is estimated to be the cause of 58,000-80,000 hospitalizations for children under the age of 5(5, 6). Recent advances in anti-RSV antivirals and vaccines for selected populations are encouraging(7–12). However, given the circulating nature of RSV among the human population, the abundance of infections, and the wide variety of clinical manifestations, there is a need for a better understanding of the genomic determinants of RSV pathogenesis. A key step in this direction is to improve the identification of RSV genotypes that are associated with different degrees of virus replication and pathogenicity.

The RSV genome is composed of 10 genes NS1, NS2, N, P, M, SH, G, F, M2, and L that encode 11 proteins. The M2 gene contains two overlapping open reading frames that code for two proteins, M2-1 and M2-2(13). The initial stage of RSV infection is determined by the virus surface glycoproteins G and F that mediate the binding of the virus to its receptor and fusion with the host cell membrane, respectively(14). G protein undergoes high selective pressure, causing frequent variations between genomes(15). The F protein, also found on the surface of the virus, is fairly conserved among different RSV strains(14, 15). Once the virus enters a cell, virus transcription, and genome replication are mediated by the polymerase L and its co-factors, including P, M2-1, and the nucleoprotein N(13, 16). M2-2 has been shown to play a role in the switch between transcription and replication, suggesting a role with the ribonucleocapsid complex(17–19).

Since its first discovery in 1956(20, 21), RSV has been classified into two main serotypes, RSV-A and RSV-B(22, 23), both of which can be further divided into multiple genotypes. Historically, due to its relatively smaller size and high variability, the evolutionary events in the G gene have been used to genotype RSV. In most cases, RSV genotypes can be distinctly defined based on the 2nd hypervariable region (HVR2) or ectodomain of G(24–29). Because G variations may impact receptor binding, G-based genotype is a good method to identify variants with different entry abilities. Whole genome sequencing has more recently been reported as an alternative method for genotyping, and it has been shown to better represent RSV genotypes in the population(30–33). M2-2 sequencing has also been proposed as an alternative genotyping method, as this is one of the smallest viral proteins with a low level of conservation(34). Notably, no current routine genotyping method focuses on the viral polymerase and its associated genes, despite the critical role of these proteins in viral replication and infection.

The lack of consensus on how to determine RSV genotypes makes it challenging to track circulating strains among the various reported sequences and even more difficult to predict associations with the replicative capacity of the different RSV variants. To assess whether variations in the replication-associated proteins can identify RSV genotypes, we took an in-depth look at the variants present in *de novo* assembled whole-length RSV genomes circulating in the US between 2012-2023. We included 31 newly sequenced samples obtained from a cohort of hospitalized children in Philadelphia, as well as 78 publicly available data sets of RSV A sequences obtained from samples around the country. Comparing RSV A sequences from samples in different US regions between the years 2012-2023, we found predictable RSV A genotypes distinguished solely by variants on the L gene. Remarkably, non-synonymous variations in L were consistently accompanied by conserved changes in the L co-factors P or M2-2, suggesting the co-variation of replication-association proteins in circulating genotypes. Furthermore, these non-synonymous variations accurately predicted RSV genotypes based on the whole genome sequence. This report demonstrates the importance of in-depth analysis of full-length RSV sequences to uncover potentially important functional changes in viral proteins, and it identifies clusters of associated variations in polymerase-related genes that can be used to genotype RSV with high accuracy.

## RESULTS

### The G and L genes are the most variable among RSV A variants circulating in the USA during 2012 – 2023

First, to identify the most variable genes in recently circulating RSV A variants, we initially sequenced and *de novo* assembled full-length genomes of 31 samples obtained from pediatric patients at the Children’s Hospital of Philadelphia (CHOP). We completely annotated these sequences and submitted them under GenBank: PP525296-PP525326. Using the consensus of the samples as a reference to identify variants between the different samples, we observed the highest number of total variations, and non-synonymous variations, in the coding regions of G and L. The M2-2 coding region showed the highest ratio of non-synonymous to total variation, and NS2 was the most conserved (Table S1). We expanded our dataset with 78 additional RSV A sequences from the National Center of Biotechnology Information (NCBI) database that accounted for different locations in the US as well as an increased range of years the sequences were collected (Table 1 and Table S2). In the combined cohort of 109 RSV A full-length sequences, we again observed the most total and non-synonymous variations in the G and L genes with the M2-2 showing the highest ratio of non-synonymous to total variation (Table 2 and Figure 1).

**Figure 1:**
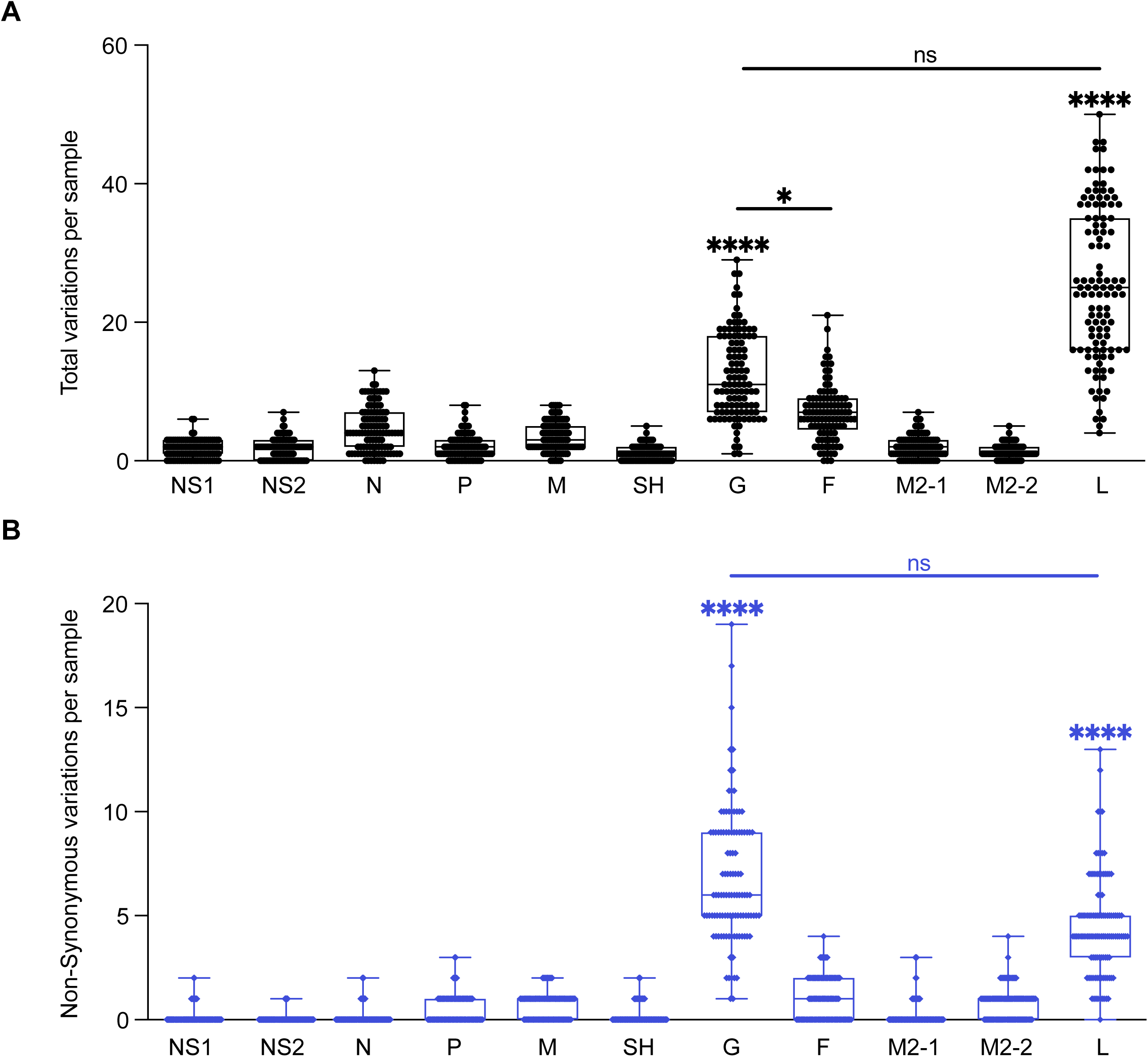
Summary of genetic variations of RSV A 11 CDS regions. Each point represents the number of variants per sample, N=109. **(A)** Total variations per sample for each CDS region. **(B)** Non-synonymous variations per sample for each CDS region. One-way analysis of variance (ANOVA) non-parametric Kruskal-Wallis tests were performed for statistical significance between CDS regions. ns p>0.1234; *p>0.03; ****p<0.0001. Comparisons between all the CDS regions to G or L were **** unless otherwise noted by a line between CDS regions.

**Table 1.** Distribution of the 109 sequences used in this study including the years samples were collected and their locations by states. All 31 PA samples are from the CHOP B Cohort and other sequences were randomly selected from NCBI. UNKN indicates that sequences are of unknown origin within the US.

|  |  | US Location |  |  |  |  |  |  |  | Total |
| --- | --- | --- | --- | --- | --- | --- | --- | --- | --- | --- |
|  |  | AR | AZ | MA | OR | PA | TN | WA | UNKN |  |
| Year | 2012 |  |  |  | 1 | 3 | 2 |  | 2 | 8 |
|  | 2013 |  |  |  |  | 7 |  |  | 1 | 8 |
|  | 2014 |  |  |  | 2 | 6 |  |  | 2 | 10 |
|  | 2015 |  |  | 1 |  | 3 |  | 4 | 3 | 11 |
|  | 2016 |  |  | 3 |  | 9 |  | 5 | 3 | 20 |
|  | 2017 |  | 1 |  |  | 1 |  |  | 2 | 4 |
|  | 2018 |  |  |  |  | 1 |  | 1 | 2 | 4 |
|  | 2019 | 1 | 2 |  |  |  |  |  | 7 | 10 |
|  | 2020 | 2 | 2 |  |  | 1 |  |  | 5 | 10 |
|  | 2021 |  |  |  |  |  |  | 1 | 3 | 4 |
|  | 2022 | 1 | 1 |  |  |  | 1 | 7 |  | 10 |
|  | 2023 | 1 | 3 |  |  |  |  | 5 | 1 | 10 |
|  | Total | 5 | 9 | 4 | 3 | 31 | 3 | 23 | 31 | 109 |

**Table 2.** Computed variations for all 109 samples showing the combined number of substitutions, insertions, and deletions per gene. Total variations were deduced from the nucleotide sequence alignment, and non-synonymous variations were deduced from the amino acid alignment.

| Gene | Total number of Variations | Number of Non-Synonymous Variations | Non-synonymous/ Total Variations |
| --- | --- | --- | --- |
| NS1 | 178 | 15 | 0.084 |
| NS2 | 190 | 6 | 0.032 |
| N | 493 | 15 | 0.030 |
| P | 220 | 56 | 0.255 |
| M | 376 | 73 | 0.194 |
| SH | 120 | 24 | 0.200 |
| G | 1348 | 743 | 0.551 |
| F | 749 | 107 | 0.143 |
| M2-1 | 219 | 27 | 0.123 |
| M2-2 | 145 | 86 | 0.593 |
| L | 2776 | 486 | 0.175 |

### Non-synonymous variations in the L gene associate with specific variations in other replication-related genes which can predict genotypic clusters

We next took a closer look at the non-synonymous variations across the coding regions of the replication-related genes (N, P, M2-1, M2-2, and L) in the 109 sequences and annotated the non-synonymous variations for each sample compared against the consensus sequence (Table S3). We found associations of variations between L and one or more of the other replication genes. We marked out the similar variations in each coding sequence (CDS) region that appeared more than two times and categorized the sequences into groups named R1-R6 based on the observed associations (Table 3, Table S4). Each group contained a varying number of sequences independent of year or location (Table S5).

**Table 3.** Definition of groups based on repetitive non-synonymous variations often seen in combination across replication-associated genes. Variations included are observed in more than 2 sequences and the groups shown account for all 109 sequences. N is not included here as there is no significant recurrence of non-synonymous variations observed. Within each CDS, a slash “/” indicates that this is another variation seen in the group. For detailed associations, see Table S4.

| Group Name | Variations in P | Variations in M2-1 | Variations in M2-2 | Variations in L |
| --- | --- | --- | --- | --- |
| R1 | I66T/ P34L |  | N46S | G1725E/ H598Y/ D1731G/ V335I/ N215S |
| R2 | T69I |  | C26Y | S1723G/ A1927T/ V1943D/ I139T/ T179A |
| R3 |  | N174S/ T180A/ S182G | L37P | N143D/ T179S/ I1653V/ K1661N/ S100L/ L953M/ S1593N/ S1789F/ H1707Q/ A2014/ Y2163N |
| R4 | T92M |  | T79A | L835M/ A1537S/ N547S/ P1720L/ V1943I |
| R5 | N59S |  |  | R511K/ R1759K |
| R6 |  |  |  | No variations |

We separated the sequences in each group based on the variation patterns observed (Table 4). R1 group contained a diverse set of variations. However, most of the R1 sequences contained L: G1725E which associated 44% of the time with M2-2: N46S. Within the sequences containing L: G1725E, three sequences had an additional L variation, N215S, which associated 100% of the time with variants, P: P34L and M2-2: N46S. Sequences with additional L variation, V335I associated 50% of the time with the P variant, I66T. The variant patterns in the R2 and R4 groups were more consistent as at least 75% of the sequences had clear associations of variations in P, M2-2, and L. Interestingly, the R3 group essentially contained L: N143D, T179S, I1653V, K1661N of which 78% did not associate with any variants in P, M2-1, or M2-2. The remaining percentage of the group did however associate with either M2-2: L37P (13%) or a trio of M2-1 variants: N174S, T180A, S182G (9%). Out of the entire cohort analyzed, R3 was the only group to consistently contain variants in M2-1. Although the R5 group had a limited number of sequences, all the sequences in this group were composed of the variants P: N59S, and L:R511K, R1759K.

**Table 4.**
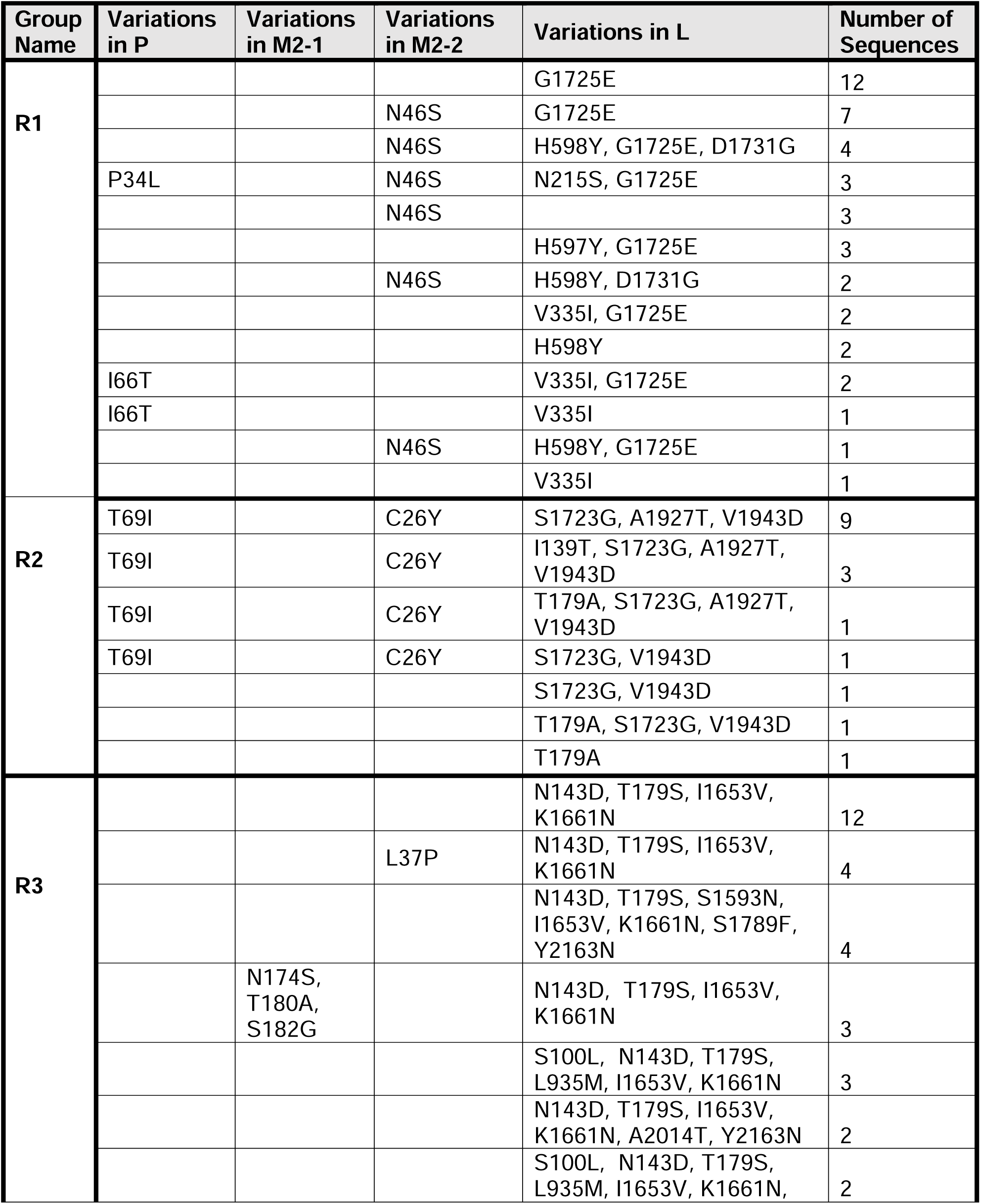
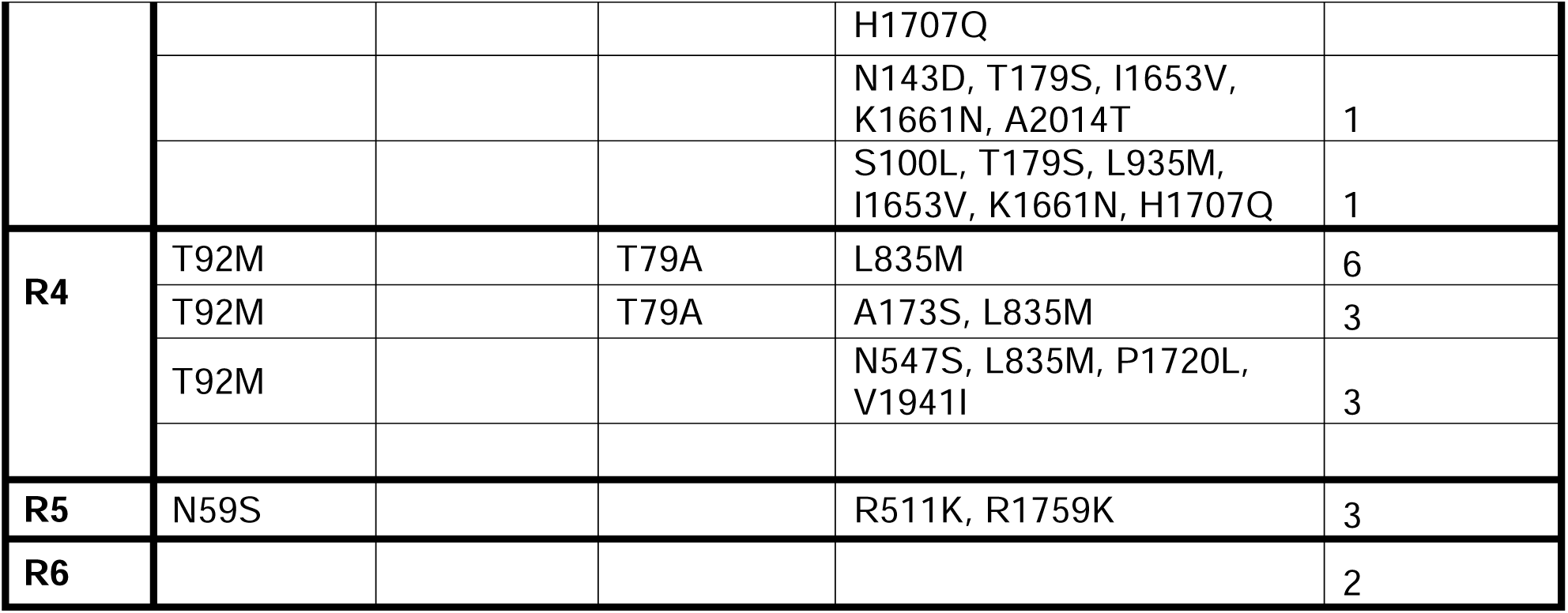
Detailed variation patterns observed in each group. N is not included here as there is no significant recurrence of non-synonymous variations observed.

Next, we generated a PCA plot with the uncorrected pairwise distances of the 109 full-length sequences and labeled them with our predicted genotypic groups R1-R6 (Figure 2A). We observed that most of the sequences distinctly clustered together based on the pre-determined groups, which supports the predicted associations among related sequences. Additionally, we labeled each sequence by the region or the year the sequences were obtained and found them to be independent of the associations (Figure 2B, C). We also employed the Nextclade web tool to assign genotypes and depict the phylogeny of the 109 samples based on 2 recently proposed classifications – Nextstrain Clades and Goya Clades(33, 35, 36). The Nextstrain Clades classification resulted in 10 clades and the Goya Clade classification resulted in only two 2 clades, GA2.3.3 and GA2.3.5 (Table S6). We labeled the previously shown PCA plot (Figure 2A) with both classification assignments from Nextclade (Figure 2D, E). Even though there are limitations to the number of clusters that can be visualized on the PCA, we found the Nextstrain Clade classification to be similar to our predicted groups. These results confirm that the non-synonymous variations in the replication-associated genes can be used to identify genotypes among RSV sequences.

**Figure 2:**
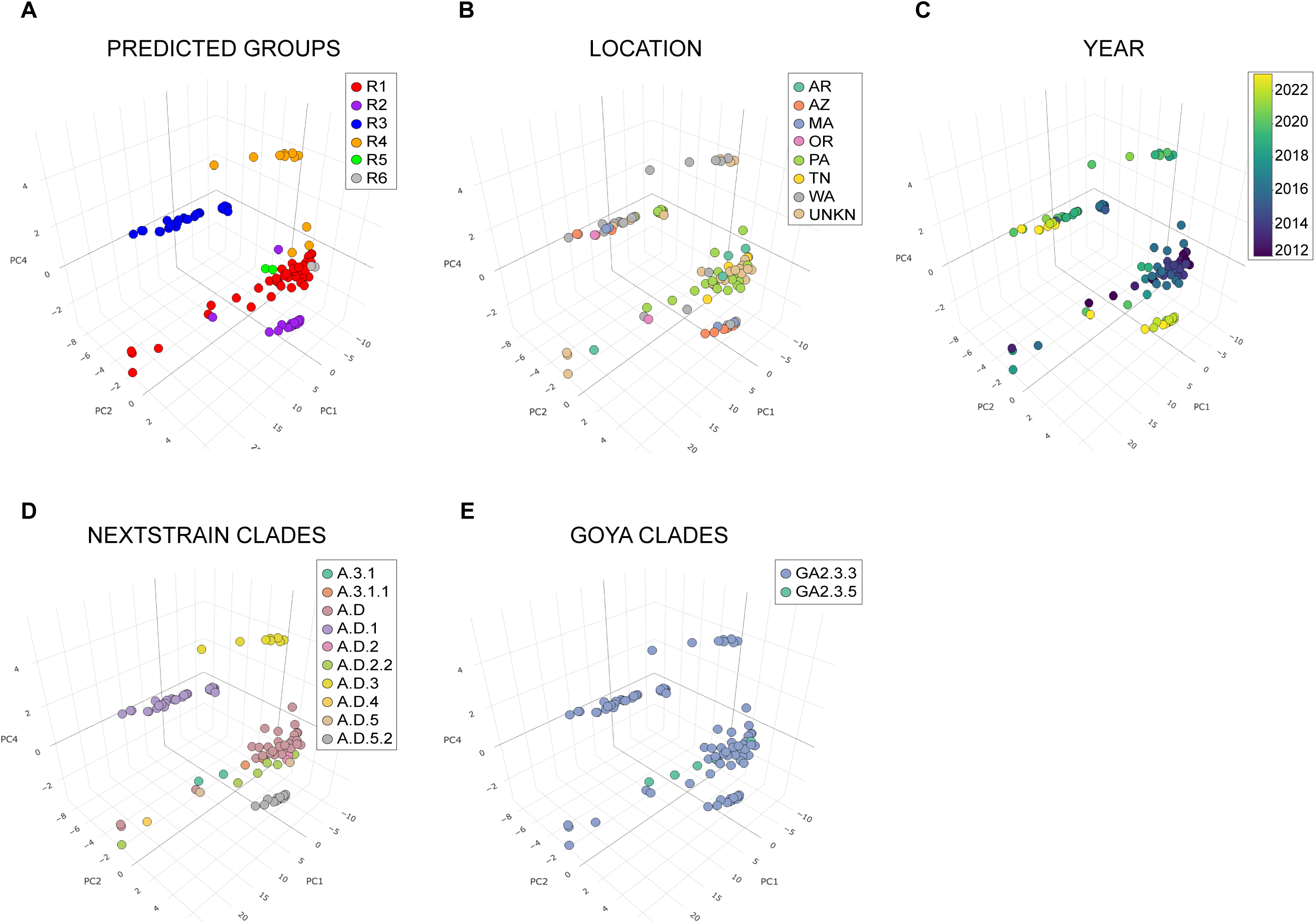
Principal component analysis (PCA) of the 109 RSV A sequences analyzed in this study. Data points were labeled based on (**A-E)** Predicted groups, Location in the US, Year the sample was collected, Nextstrain Clades classification, and Goya Clades classification respectively. The percentage of variance in all the plots are PC1=58.71%, PC2=21.78%, and PC4=2.52%.

### Non-synonymous variations in replication-associated genes concentrate in protein-protein interaction domains

To determine if specific domains of the RSV replication-associated proteins were susceptible to variation, we annotated the variants observed in the different groups to their respective predicted domain residues (Figure 3)(16). All the variants observed in P were found in the N-terminal domain (NTD). The NTD of P is thought to bind to M2-1, specifically regions of the core domain, as well as interact with RNA-free N monomers(16). The variants of M2-1 were limited to the core domain. The structure of M2-2 has yet to be resolved, therefore no domains are assigned to the protein.

**Figure 3:**
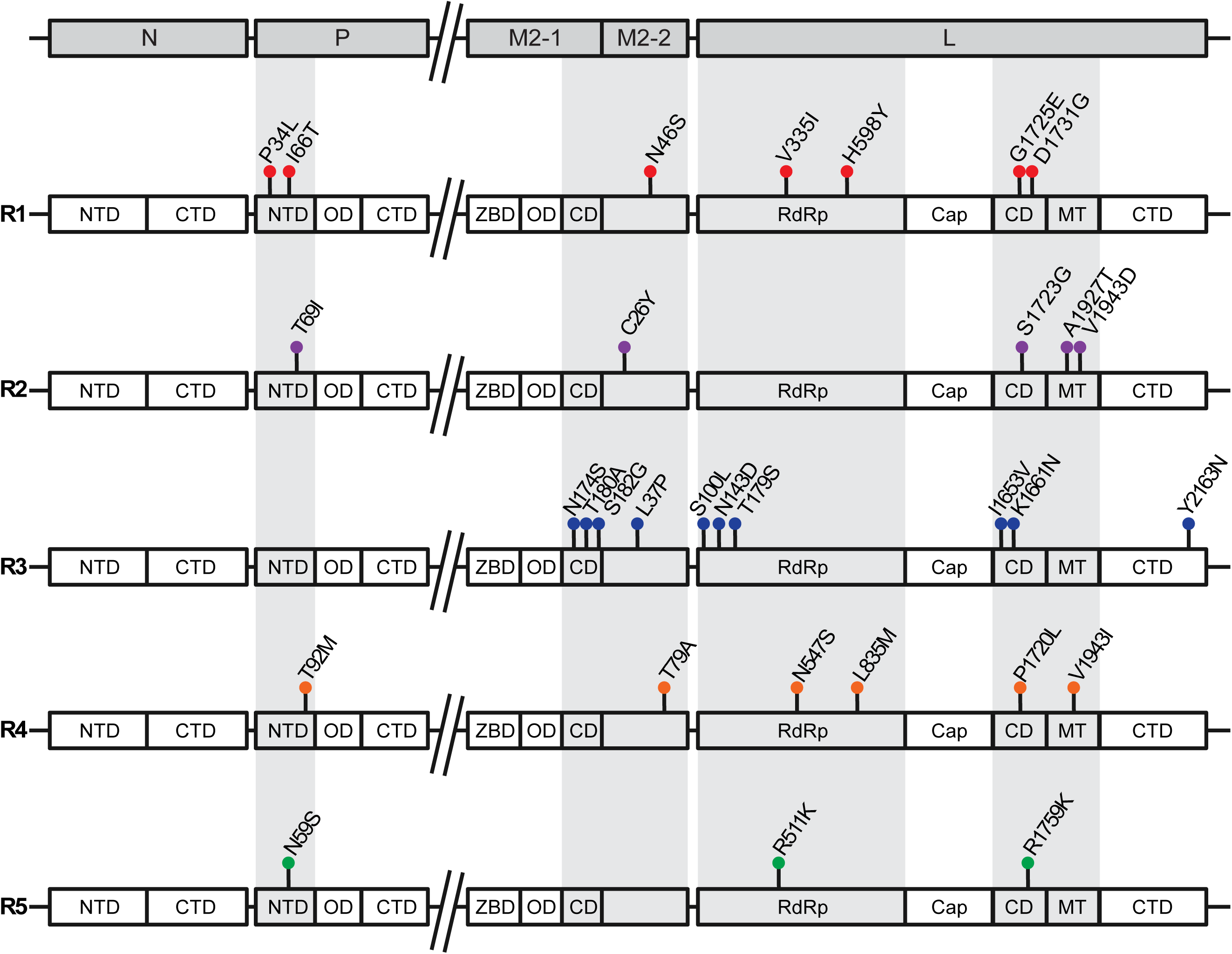
Annotation of non-synonymous variations observed within the domains RSV A replication-associated genes for each predicted group. All variations shown are seen at least 3 times and interact with other variations within a predicted group. Abbreviation key as follows: NTD: N-terminal domain; CTD: C-terminal domain; OD: oligomerization domain; ZBD: Zinc-binding domain; M2-1 CD: core domain; RdRP: RNA-dependent RNA polymerase; Cap: cap addition domain; L CD: connector domain; MT: cap methylation domain.

However, it has been shown that overexpression of M2-2 rearranges the ribonucleocapsid complex, suggesting potential interactions with L(19). We observed most variants of L in the RNA-dependent RNA-polymerase (RdRp), connector domain (CD), and methyltransferase (MT) domains. It has been shown that the RdRp of L is an interaction site for P(37, 38). The complete structures of the CD and MT domains have not been determined in the attempts to resolve the full L structure; however, it is suggested that these domains may be dynamic and also interact with the P protein(16, 37, 38). Overall, these associations indicate potential residues of interest in the replication-associated proteins that may impact the fitness of the virus during an infection.

## DISCUSSION

Our in-depth analysis of RSV A sequences shows distinct clusters of variations within replication-associated genes as specific variation(s) in L often associate with variations in one or more of the other replication-associated genes. We grouped these associations, depicted them in a PCA plot, and compared their similarity to relatively known methods of RSV genotyping with the Nextclade tool. Our observed associations were not only capable of defining genotypic groups, but also support the advantage of full-length sequence analyses to properly characterize and genotype RSV subgroups. Importantly, our analysis revealed associations of variants among different replication-associated genes. These data can be used to infer the dynamic variation of RSV genotypes and their associated pathogenicity in different seasons, as well as as starting point for detailed investigation of the molecular interactions among proteins from the replication complex.

Most of the reported sequencing analyses from clinical studies generate new sequences based on a known or closely related reference sequence(39–42). An alternative method is *de novo* assembly. The advantage of *de novo* assembly is that it does not require a reference sequence that could potentially introduce bias in the form of variations in the consensus sequence. A merger known as reference-guided *de novo* assembly has also been proposed for larger genome construction (43). In this study, we employed the SPAdes *de novo* assembler with rnaviralSPAdes option which is well adapted for shorter reads and metavirome (44, 45). This gives an accurate sequence per isolate as genomic variations including intergenic, or repetitive regions, and indels are also accounted for.

With *de novo* assembly one must be conscientious when trimming the CDS regions. For RSV, one of the prominent characteristics observed during alignment is the novel duplication in the C-terminal region of the attachment glycoprotein(25). Of the 109 sequences analyzed in this study, only 10 were without the duplication (GenBank: KY967362, MF001039, PP525326, MN306050, MN306054, PP525323, OK659680, PP525325, PP5253324, MN310477). In addition to the duplication event in the G, we also observed the presence of alternate codons while trimming the CDS regions. We observed that these alternate codons often lead to a few extra or lesser amino acidd that may not be properly annotated during database submissions. Observed alternate codons include 7 sequences that have a later stop codon in the G gene (GenBank: OK649680-84, 0R287918, OR287948). This alternate codon is also present in the well-studied reference genome – GenBank: KT992094.1. Two sequences were also observed to have a later stop codon in the L gene in this study, which has not been previously reported (GenBank: PP525304, OR143185). Other sequences without these later stop codons often have a tandem stop codon e.g. “TAATGA”. These tandem stop codons have been studied in the RSV G gene and shown to decrease the expression of fusion glycoprotein (F) and facilitate immune evasion of the virus resulting in the severity of the infection(46, 47). Nine sequences in this study have an alternative start codon in the M2-2 gene (GenBank: PP525314, PP525323-4, OK649682, OK649684, OR143171, OR143187, OR522529, OR601479). M2-2 is generally known to have three start codons and HRSV A2 has this variation too(18, 48). Only the first and second of these start codons have been shown to result in a functional M2-2 protein during replication(18). Alternate start codons in the Influenza virus have been shown not to affect viral fitness, particularly when the translated protein does not result in a missing domain(49, 50).

One unclear question is which protein(s) is/are the major driver of genomic variations: the surface proteins – SH, G, and F that are subjected to host immune activities or the polymerase protein L responsible for ensuring viral growth fitness. The answer could be both. It has been established that the 2nd hypervariable region of G (HVR2) is highly variable and often used in determining RSV genotypes(51). However, in this study, we focused on a replication aspect starting with the polymerase L and observed variations majorly in the RdRP, CD, and MT domains. Consistent associations were also observed in other replication-associated genes such as the NTD of P, the core domain of M2-1, and M2-2. It is known that L, but not P, interacts directly with the RNA template and P interacts with L as a cofactor during replication (52). Established protein prediction also shows that L binds to the CTD of P rather than its NTD(16, 38). However, in the well-studied Vesicular Stomatitis Virus (VSV), it has been shown that the NTD of P interacts with the CTD, RdRp, and CD domains of L(53–55). The CD, MT, and CTD in VSV L have no fixed position and it is this compact conformation with P that transforms it to an initiation competent state(53). With such an interactive polymerase complex there are multiple ways the variations observed may impact the functionality of the virus, such as speed or fidelity. These factors should also be considered when tracking and tracing different RSV genotypes.

With the prevalence of circulating RSV among the human population, it is crucial to understand the genomic determinants of RSV pathogenesis. It has been shown that different RSV strains may impact clinical outcomes(47, 56). Yet, there is a lack of consensus on the exact strains that cause the different severities of disease (57).

Improving the annotation of variations specifically involved in virus replication may aid in the identification of RSV genotypes that correspond to pathogenesis. From our results, we support that full-length sequencing is the most reliable data required for genotyping RSV A, closely followed by genotyping the L gene. We also suggest investigating the variation patterns in the CDS of replication-associated genes can be very informative in downstream studies on the efficiency of the polymerase.

## MATERIALS AND METHODS

### Description of cohort and Data collection

Ninety-six RSV clinical samples were obtained from the Children’s Hospital of Philadelphia (CHOP), Philadelphia, Pennsylvania, USA. Samples used in this study were from 3 different coded cohorts. Cohort H (7 samples; 2012 and 2017), Cohort CL (6 samples; 2013 – 2015), and Cohort B (83 samples; 2015 and 2016). All clinical samples were nasal washes collected from hospitalized patients between the age of 0-2 years as reviewed and approved by the CHOP Institutional Review Board (IRB). From the 96 samples, we could *de novo* assemble full-length sequences for the 31 included in this study. Other sample sequences analyzed outside these cohorts were obtained from the National Center for Biotechnology Information (NCBI Virus) and Bacterial and Viral Bioinformatics Resource Center (BV-BRC) databases (78 sequences; 2012 - 2023).

Complete sequences without insertions in their CDS were selected at random to increase the number of sequences we have each year (Table S2).

### RNA extraction, library preparation, and NGS sequencing

The processing of samples in Cohort H (7 samples) and CL (6 samples) has been previously described in detail(58, 59). For Cohort B (83 samples), total RNA was extracted from clinical samples using TRIzol LS (Invitrogen) according to the manufacturer’s instructions. Linear Acrylamide (Invitrogen) was added at the precipitation step of RNA extraction to increase yield. RNA quantity and quality were measured using Nanodrop and Bioanalyzer (Agilent Technologies).

Sigma SeqPlex RNA Amplification Kit was used for making the complementary deoxyribonucleic acid (cDNA) library preparation for all samples to be sequenced using the Illumina NovaSeq 6000 to generate 150-bp, paired-end reads. The sequencing generated an average of 50 million reads per sample with an average Phred quality score between 34.6 and 36.1.

### De novo assembly

Illumina adaptors were removed from the resulting Illumina paired-end reads using Cutadapt (v2.5)(60) and the trimmed reads were analyzed using FastQC (v0.11) to ensure their quality. Bowtie2 (v2.4.1)(61) was then used to remove trimmed reads that aligned to the human genome (GRCh38) to generate non-host reads for subsequent analysis. St. Petersburg genome assembler (SPAdes) v3.15.5(44) was used for the de novo assembly of the non-host paired-end reads of each sample. An additional pipeline option in the assembler recommended for RNA viral datasets (-- rnaviral) and a thread of 50 were included as parameters while running the assembler with docker on the command line. The contigs and scaffolds generated by the assembler for each sample were aligned against Genbank: KC731482.1 using BLASTN to identify and obtain the RSV A sequences used in this study.

### Variant analysis

MegAlign Pro v17.5.0 (DNASTAR) was used for most of the analysis of the selected sequences of RSV A in this study. The complete nucleotide sequences of all samples were aligned with GenBank: KC731482.1, KT992094.1 using MUSCLE with default options. The coding sequences (CDS) of RSV A (NS1, NS2, N, M, P, G, F, SH, M2-1, M2-2, and L) were obtained by trimming the aligned sequences according to the annotation of the references from their feature tables on GenBank. EMBOSS Transeq tool (EMBL-EBI)(62) was used to translate the trimmed CDS to amino acids. The resulting amino acid sequences for each gene were re-uploaded and re-aligned with MUSCLE, and the reference sequences used in trimming the CDS were excluded from subsequent analysis. Variant calling for each CDS was generated using “Compute Variants” in MegAlign Pro v17.5.0 after which these were compiled as reference amino acid, (X) reference position, (123) variant in the sample (Y) i.e. X123Y. The reference used in this analysis is the consensus sequence generated after the alignment of all the sequences.

### PCA

Principal component analysis graphs were plotted in R Studio v 2023.12.1+402 using plotly library. The input data were the demographics of the sequences and their uncorrected pairwise distance matrix.

### Clades assignment

Assignment of Nextstrain and Goya clades were made in Nextclade v3.2.0 https://clades.nextstrain.org(36).

## DATA AVAILABILITY

All raw sequencing data used for *de novo* assembly were deposited in SRA under accession numbers PRJNA837014, and PRJNA681672. Assembled and annotated sequences were submitted in GenBank and were assigned accession numbers PP525296-PP525326.

## ACKNOWLEDGEMENTS

This work was supported by the U.S. National Institutes of Health National Institute of Allergy and Infectious Diseases (R01 AI137062 to CLB) and the BJC Investigators Program. The authors wish to acknowledge Emna Achouri for training and support with Bioinformatics analyses and Matt Devine for help with the data pull and identification of CHOP samples.

## SUPPLEMENTARY TABLES AND LEGENDS

**Table S1:**
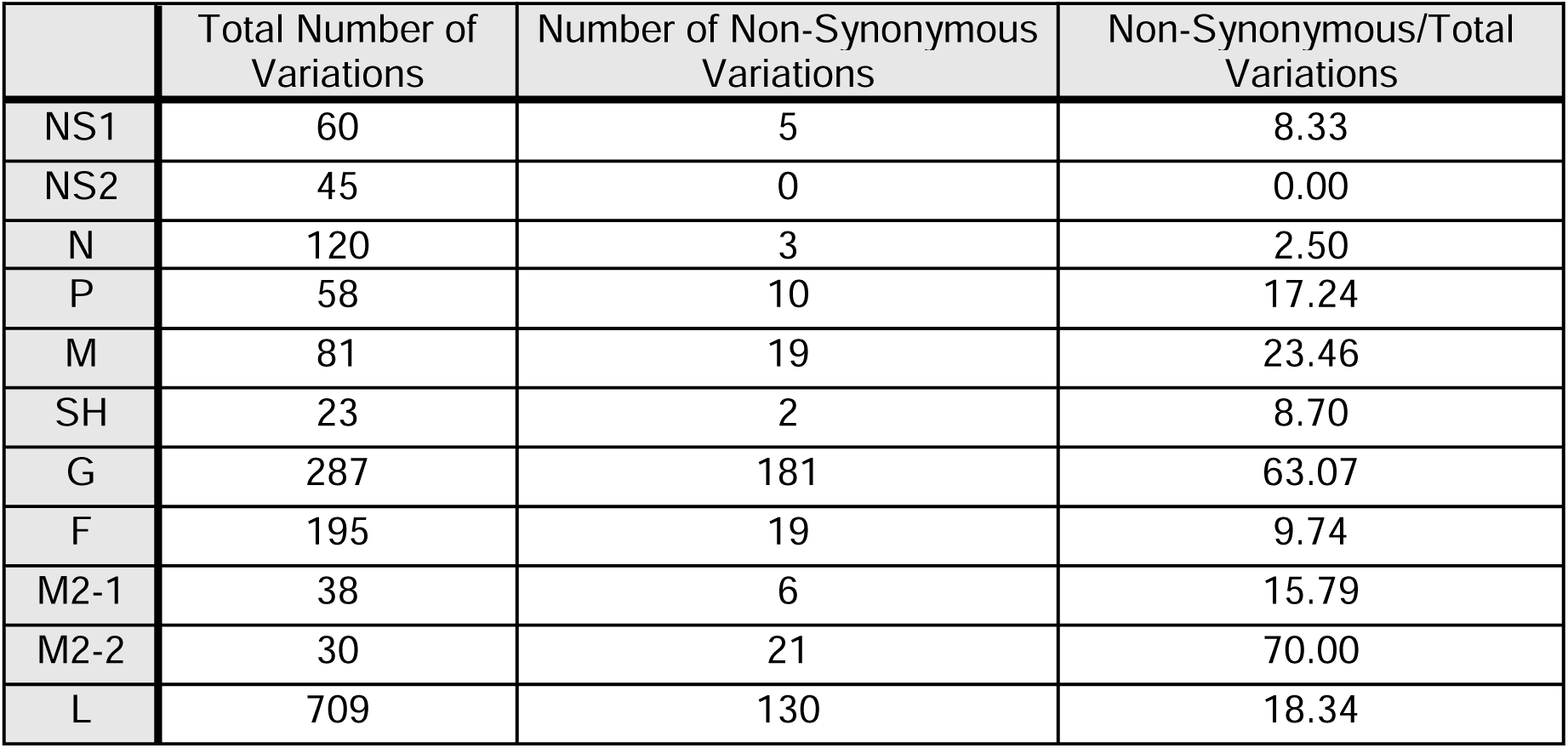
Variations in the 31 RSV A sequences from Philadelphia. Computed variations showing the combined number of substitutions, insertions, and deletions per gene. Total variations were deduced from the nucleotide sequence alignment, and non- synonymous variations were deduced from the amino acid alignment.

**Table S2:**
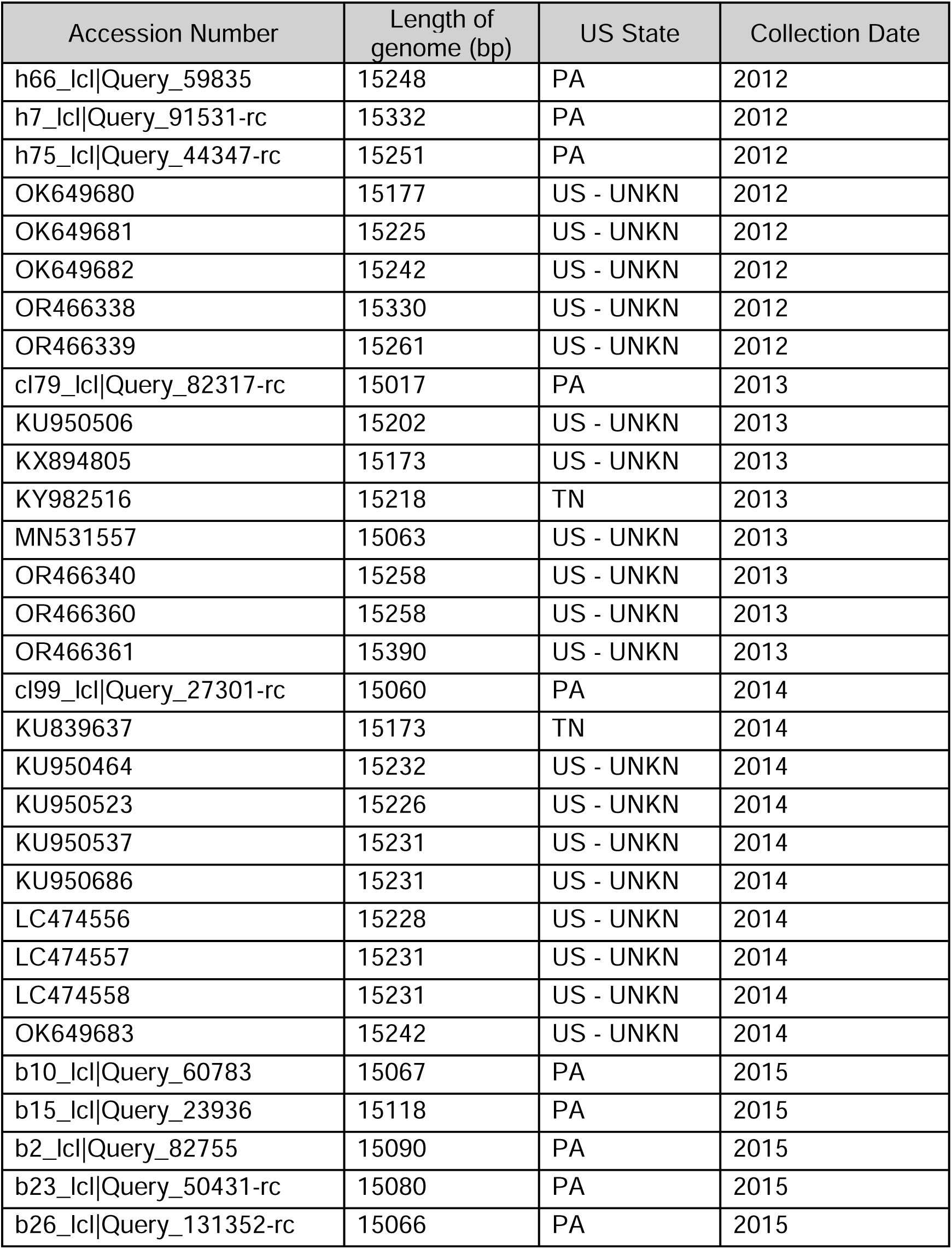
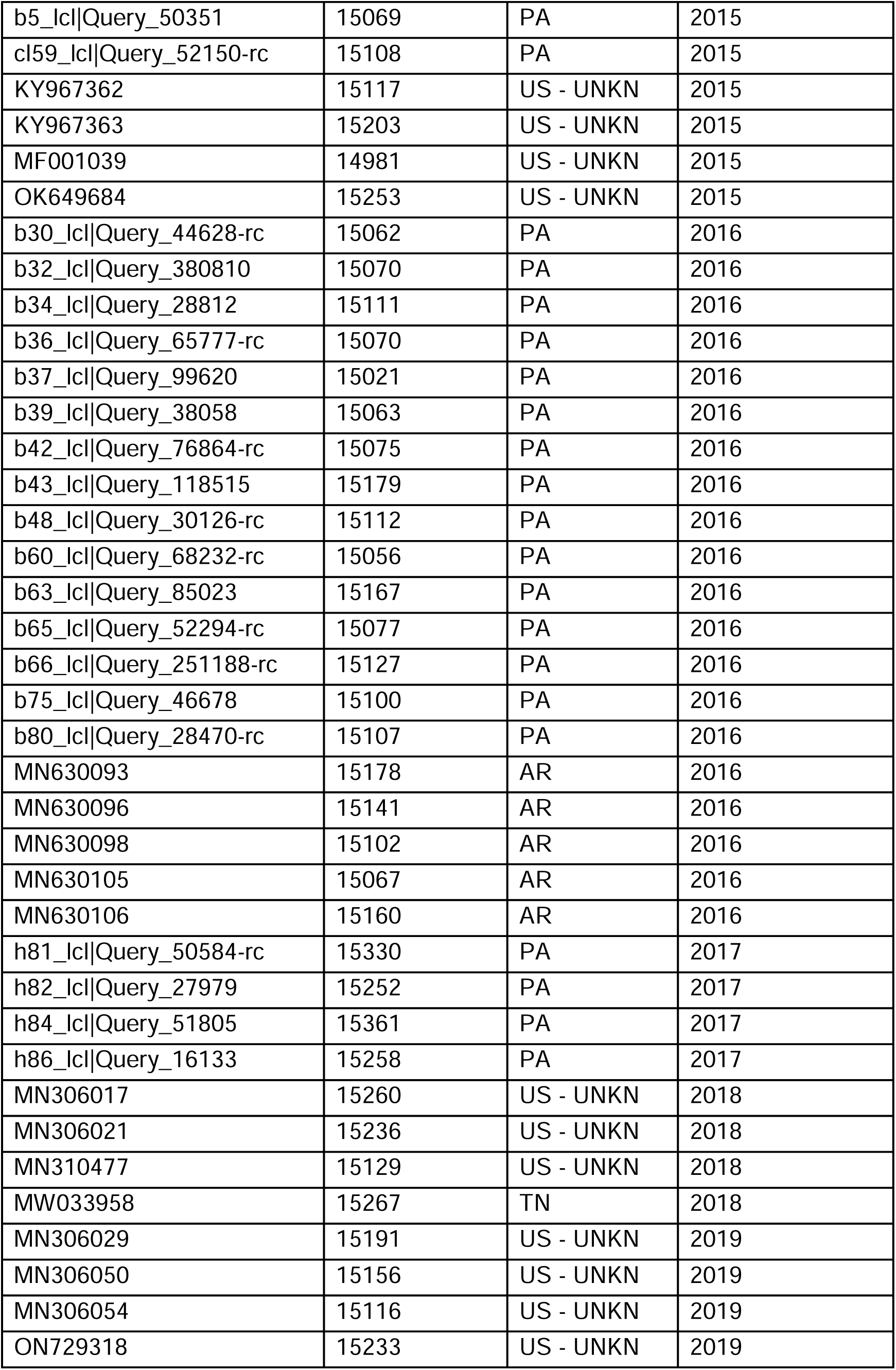
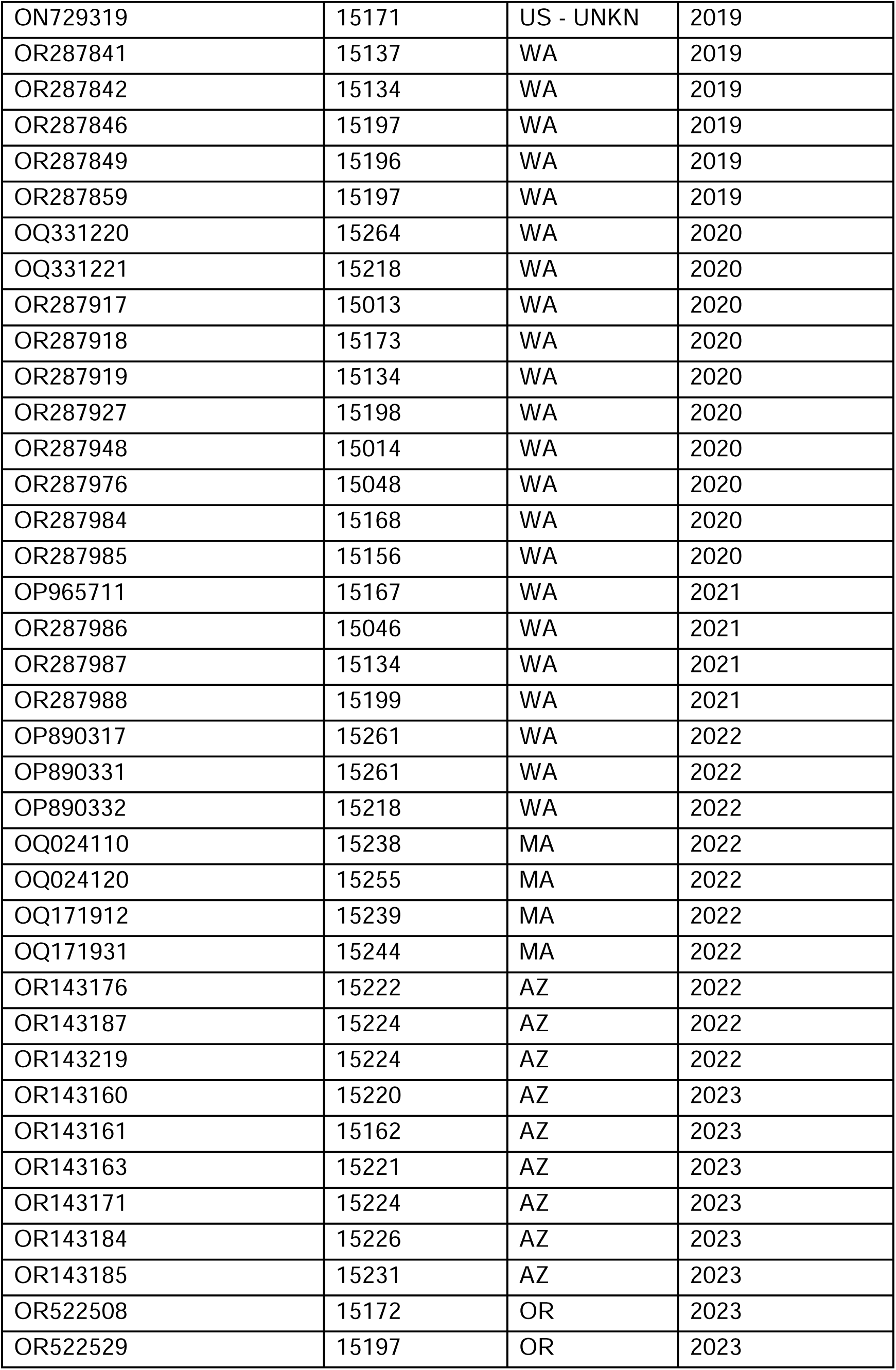
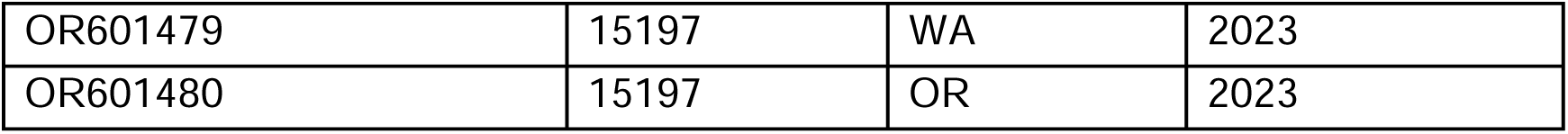
GenBank accession number, length of genome, US states, year of sample collection of all 109 RSV A sequences. UNKN indicates sequences of unknown origin within the US.

| Accession Number | Length of genome (bp) | US State | Collection Date |
| --- | --- | --- | --- |
| h66_lcl\|Query_59835 | 15248 | PA | 2012 |
| h7_lcl\|Query_91531-rc | 15332 | PA | 2012 |
| h75_lcl\|Query_44347-rc | 15251 | PA | 2012 |
| OK649680 | 15177 | US - UNKN | 2012 |
| OK649681 | 15225 | US - UNKN | 2012 |
| OK649682 | 15242 | US - UNKN | 2012 |
| OR466338 | 15330 | US - UNKN | 2012 |
| OR466339 | 15261 | US - UNKN | 2012 |
| cl79_lcl\|Query_82317-rc | 15017 | PA | 2013 |
| KU950506 | 15202 | US - UNKN | 2013 |
| KX894805 | 15173 | US - UNKN | 2013 |
| KY982516 | 15218 | TN | 2013 |
| MN531557 | 15063 | US - UNKN | 2013 |
| OR466340 | 15258 | US - UNKN | 2013 |
| OR466360 | 15258 | US - UNKN | 2013 |
| OR466361 | 15390 | US - UNKN | 2013 |
| cl99_lcl\|Query_27301-rc | 15060 | PA | 2014 |
| KU839637 | 15173 | TN | 2014 |
| KU950464 | 15232 | US - UNKN | 2014 |
| KU950523 | 15226 | US - UNKN | 2014 |
| KU950537 | 15231 | US - UNKN | 2014 |
| KU950686 | 15231 | US - UNKN | 2014 |
| LC474556 | 15228 | US - UNKN | 2014 |
| LC474557 | 15231 | US - UNKN | 2014 |
| LC474558 | 15231 | US - UNKN | 2014 |
| OK649683 | 15242 | US - UNKN | 2014 |
| b10_lcl\|Query_60783 | 15067 | PA | 2015 |
| b15_lcl\|Query_23936 | 15118 | PA | 2015 |
| b2_lcl\|Query_82755 | 15090 | PA | 2015 |
| b23_lcl\|Query_50431-rc | 15080 | PA | 2015 |
| b26_lcl\|Query_131352-rc | 15066 | PA | 2015 |
| b5_lcl\|Query_50351 | 15069 | PA | 2015 |
| cl59_lcl\|Query_52150-rc | 15108 | PA | 2015 |
| KY967362 | 15117 | US - UNKN | 2015 |
| KY967363 | 15203 | US - UNKN | 2015 |
| MF001039 | 14981 | US - UNKN | 2015 |
| OK649684 | 15253 | US - UNKN | 2015 |
| b30_lcl\|Query_44628-rc | 15062 | PA | 2016 |
| b32_lcl\|Query_380810 | 15070 | PA | 2016 |
| b34_lcl\|Query_28812 | 15111 | PA | 2016 |
| b36_lcl\|Query_65777-rc | 15070 | PA | 2016 |
| b37_lcl\|Query_99620 | 15021 | PA | 2016 |
| b39_lcl\|Query_38058 | 15063 | PA | 2016 |
| b42_lcl\|Query_76864-rc | 15075 | PA | 2016 |
| b43_lcl\|Query_118515 | 15179 | PA | 2016 |
| b48_lcl\|Query_30126-rc | 15112 | PA | 2016 |
| b60_lcl\|Query_68232-rc | 15056 | PA | 2016 |
| b63_lcl\|Query_85023 | 15167 | PA | 2016 |
| b65_lcl\|Query_52294-rc | 15077 | PA | 2016 |
| b66_lcl\|Query_251188-rc | 15127 | PA | 2016 |
| b75_lcl\|Query_46678 | 15100 | PA | 2016 |
| b80_lcl\|Query_28470-rc | 15107 | PA | 2016 |
| MN630093 | 15178 | AR | 2016 |
| MN630096 | 15141 | AR | 2016 |
| MN630098 | 15102 | AR | 2016 |
| MN630105 | 15067 | AR | 2016 |
| MN630106 | 15160 | AR | 2016 |
| h81_lcl\|Query_50584-rc | 15330 | PA | 2017 |
| h82_lcl\|Query_27979 | 15252 | PA | 2017 |
| h84_lcl\|Query_51805 | 15361 | PA | 2017 |
| h86_lcl\|Query_16133 | 15258 | PA | 2017 |
| MN306017 | 15260 | US - UNKN | 2018 |
| MN306021 | 15236 | US - UNKN | 2018 |
| MN310477 | 15129 | US - UNKN | 2018 |
| MW033958 | 15267 | TN | 2018 |
| MN306029 | 15191 | US - UNKN | 2019 |
| MN306050 | 15156 | US - UNKN | 2019 |
| MN306054 | 15116 | US - UNKN | 2019 |
| ON729318 | 15233 | US - UNKN | 2019 |
| ON729319 | 15171 | US - UNKN | 2019 |
| OR287841 | 15137 | WA | 2019 |
| OR287842 | 15134 | WA | 2019 |
| OR287846 | 15197 | WA | 2019 |
| OR287849 | 15196 | WA | 2019 |
| OR287859 | 15197 | WA | 2019 |
| OQ331220 | 15264 | WA | 2020 |
| OQ331221 | 15218 | WA | 2020 |
| OR287917 | 15013 | WA | 2020 |
| OR287918 | 15173 | WA | 2020 |
| OR287919 | 15134 | WA | 2020 |
| OR287927 | 15198 | WA | 2020 |
| OR287948 | 15014 | WA | 2020 |
| OR287976 | 15048 | WA | 2020 |
| OR287984 | 15168 | WA | 2020 |
| OR287985 | 15156 | WA | 2020 |
| OP965711 | 15167 | WA | 2021 |
| OR287986 | 15046 | WA | 2021 |
| OR287987 | 15134 | WA | 2021 |
| OR287988 | 15199 | WA | 2021 |
| OP890317 | 15261 | WA | 2022 |
| OP890331 | 15261 | WA | 2022 |
| OP890332 | 15218 | WA | 2022 |
| OQ024110 | 15238 | MA | 2022 |
| OQ024120 | 15255 | MA | 2022 |
| OQ171912 | 15239 | MA | 2022 |
| OQ171931 | 15244 | MA | 2022 |
| OR143176 | 15222 | AZ | 2022 |
| OR143187 | 15224 | AZ | 2022 |
| OR143219 | 15224 | AZ | 2022 |
| OR143160 | 15220 | AZ | 2023 |
| OR143161 | 15162 | AZ | 2023 |
| OR143163 | 15221 | AZ | 2023 |
| OR143171 | 15224 | AZ | 2023 |
| OR143184 | 15226 | AZ | 2023 |
| OR143185 | 15231 | AZ | 2023 |
| OR522508 | 15172 | OR | 2023 |
| OR522529 | 15197 | OR | 2023 |
| OR601479 | 15197 | WA | 2023 |
| OR601480 | 15197 | OR | 2023 |

**Table S3:** Annotated variations observed in the CDS of replication-associated genes (N, P, M2, L) of selected 109 RSV A sequences. An asterisk “*” indicates that the sequence has no variation in the gene when compared to the consensus sequence. **See Excel file uploaded**.

**Table S4:** Complete version of Table 4 showing variations observed in more than 2 sequences. Variations are arranged by their positions in the CDS and assigned to one of the groups R1-R6. **See Excel file uploaded**.

**Table S5:**
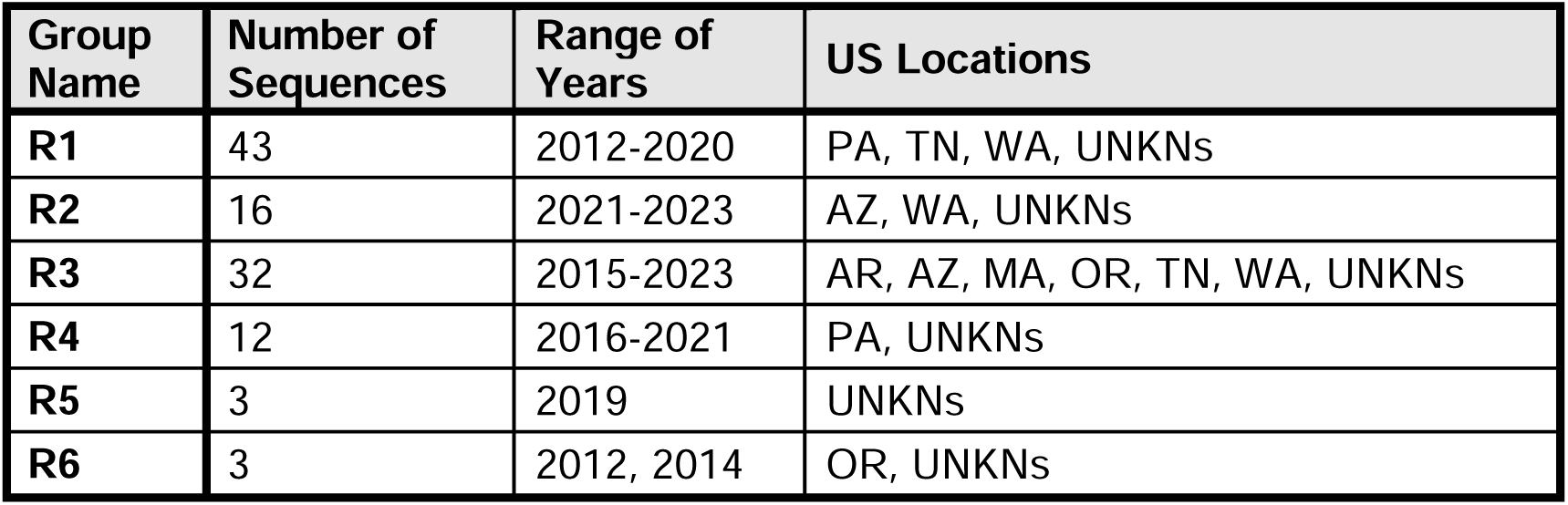
Distribution of 109 sequences within each predicted groups including the year of sample collection and their locations in the US by states. UNKNs indicates that sequences are of unknown origin within the US.

**Table S6:** Table showing assigned Nextstrain clade and Goya clade of each sequence compared to our predicted R1-R6 groups. Clades were determined using full-length sequences in the Nextclade tool. **See Excel file uploaded**.

## REFERENCES

1. Glezen WP, Taber LH, Frank AL, Kasel JA. 1986. Risk of primary infection and reinfection with respiratory syncytial virus. Am J Dis Child 140:543–6.

2. Collins PL, Melero JA. 2011. Progress in understanding and controlling respiratory syncytial virus: still crazy after all these years. Virus Res 162:80–99.

3. Pneumonia Etiology Research for Child Health Study G. 2019. Causes of severe pneumonia requiring hospital admission in children without HIV infection from Africa and Asia: the PERCH multi-country case-control study. Lancet 394:757- 779.

4. Falsey AR, Hennessey PA, Formica MA, Cox C, Walsh EE. 2005. Respiratory syncytial virus infection in elderly and high-risk adults. N Engl J Med 352:1749–59.

5. Hall CB, Weinberg GA, Iwane MK, Blumkin AK, Edwards KM, Staat MA, Auinger P, Griffin MR, Poehling KA, Erdman D, Grijalva CG, Zhu Y, Szilagyi P. 2009. The burden of respiratory syncytial virus infection in young children. N Engl J Med 360:588–98.

6. McLaughlin JM, Khan F, Schmitt HJ, Agosti Y, Jodar L, Simoes EAF, Swerdlow DL. 2022. Respiratory Syncytial Virus-Associated Hospitalization Rates among US Infants: A Systematic Review and Meta-Analysis. Journal of Infectious Diseases 225:1100–1111.

7. Griffin MP, Yuan Y, Takas T, Domachowske JB, Madhi SA, Manzoni P, Simoes EAF, Esser MT, Khan AA, Dubovsky F, Villafana T, DeVincenzo JP, Nirsevimab Study G. 2020. Single-Dose Nirsevimab for Prevention of RSV in Preterm Infants. N Engl J Med 383:415–425.

8. Jones JM, Fleming-Dutra KE, Prill MM, Roper LE, Brooks O, Sanchez PJ, Kotton CN, Mahon BE, Meyer S, Long SS, Mcmorrow ML. 2023. Use of Nirsevimab for the Prevention of Respiratory Syncytial Virus Disease Among Infants and Young Children: Recommendations of the Advisory Committee on Immunization Practices - United States, 2023. Mmwr-Morbidity and Mortality Weekly Report 72:920-925.

9. Papi A, Ison MG, Langley JM, Lee DG, Leroux-Roels I, Martinon-Torres F, Schwarz TF, van Zyl-Smit RN, Campora L, Dezutter N, de Schrevel N, Fissette L, David MP, Van der Wielen M, Kostanyan L, Hulstrom V, Group AR-S. 2023. Respiratory Syncytial Virus Prefusion F Protein Vaccine in Older Adults. N Engl J Med 388:595–608.

10. Walsh EE, Perez Marc G, Zareba AM, Falsey AR, Jiang Q, Patton M, Polack FP, Llapur C, Doreski PA, Ilangovan K, Ramet M, Fukushima Y, Hussen N, Bont LJ, Cardona J, DeHaan E, Castillo Villa G, Ingilizova M, Eiras D, Mikati T, Shah RN, Schneider K, Cooper D, Koury K, Lino MM, Anderson AS, Jansen KU, Swanson KA, Gurtman A, Gruber WC, Schmoele-Thoma B, Group RCT. 2023. Efficacy and Safety of a Bivalent RSV Prefusion F Vaccine in Older Adults. N Engl J Med 388:1465–1477.

11. Melgar M, Britton A, Roper LE, Talbot HK, Long SS, Kotton CN, Havers FP. 2023. Use of Respiratory Syncytial Virus Vaccines in Older Adults: Recommendations of the Advisory Committee on Immunization Practices - United States, 2023. MMWR Morb Mortal Wkly Rep 72:793-801.

12. Kampmann B, Madhi SA, Munjal I, Simoes EAF, Pahud BA, Llapur C, Baker J, Perez Marc G, Radley D, Shittu E, Glanternik J, Snaggs H, Baber J, Zachariah P, Barnabas SL, Fausett M, Adam T, Perreras N, Van Houten MA, Kantele A, Huang LM, Bont LJ, Otsuki T, Vargas SL, Gullam J, Tapiero B, Stein RT, Polack FP, Zar HJ, Staerke NB, Duron Padilla M, Richmond PC, Koury K, Schneider K, Kalinina EV, Cooper D, Jansen KU, Anderson AS, Swanson KA, Gruber WC, Gurtman A, Group MS. 2023. Bivalent Prefusion F Vaccine in Pregnancy to Prevent RSV Illness in Infants. N Engl J Med 388:1451–1464.

13. Collins PL, Fearns R, Graham BS. 2013. Respiratory syncytial virus: virology, reverse genetics, and pathogenesis of disease. Curr Top Microbiol Immunol 372:3–38.

14. McLellan JS, Ray WC, Peeples ME. 2013. Structure and function of respiratory syncytial virus surface glycoproteins. Curr Top Microbiol Immunol 372:83–104.

15. Tan L, Coenjaerts FE, Houspie L, Viveen MC, van Bleek GM, Wiertz EJ, Martin DP, Lemey P. 2013. The comparative genomics of human respiratory syncytial virus subgroups A and B: genetic variability and molecular evolutionary dynamics. J Virol 87:8213–26.

16. Cao D, Gao Y, Liang B. 2021. Structural Insights into the Respiratory Syncytial Virus RNA Synthesis Complexes. Viruses 13.

17. Bermingham A, Collins PL. 1999. The M2-2 protein of human respiratory syncytial virus is a regulatory factor involved in the balance between RNA replication and transcription. Proc Natl Acad Sci U S A 96:11259–64.

18. Cheng X, Park H, Zhou H, Jin H. 2005. Overexpression of the M2-2 protein of respiratory syncytial virus inhibits viral replication. J Virol 79:13943–52.

19. Blanchard EL, Braun MR, Lifland AW, Ludeke B, Noton SL, Vanover D, Zurla C, Fearns R, Santangelo PJ. 2020. Polymerase-tagged respiratory syncytial virus reveals a dynamic rearrangement of the ribonucleocapsid complex during infection. PLoS Pathog 16:e1008987.

20. Blount RE, Jr., Morris JA, Savage RE. 1956. Recovery of cytopathogenic agent from chimpanzees with coryza. Proc Soc Exp Biol Med 92:544–9.

21. Chanock R, Roizman B, Myers R. 1957. Recovery from infants with respiratory illness of a virus related to chimpanzee coryza agent (CCA). I. Isolation, properties and characterization. Am J Hyg 66:281–90.

22. Mufson MA, Orvell C, Rafnar B, Norrby E. 1985. Two distinct subtypes of human respiratory syncytial virus. J Gen Virol 66 ( Pt 10):2111–24.

23. Anderson LJ, Hierholzer JC, Tsou C, Hendry RM, Fernie BF, Stone Y, McIntosh K. 1985. Antigenic characterization of respiratory syncytial virus strains with monoclonal antibodies. J Infect Dis 151:626–33.

24. Sullender WM. 2000. Respiratory syncytial virus genetic and antigenic diversity. Clin Microbiol Rev 13:1–15, table of contents.

25. Eshaghi A, Duvvuri VR, Lai R, Nadarajah JT, Li A, Patel SN, Low DE, Gubbay JB. 2012. Genetic variability of human respiratory syncytial virus A strains circulating in Ontario: a novel genotype with a 72 nucleotide G gene duplication. PLoS One 7:e32807.

26. Agoti CN, Otieno JR, Gitahi CW, Cane PA, Nokes DJ. 2014. Rapid spread and diversification of respiratory syncytial virus genotype ON1, Kenya. Emerg Infect Dis 20:950–9.

27. Munoz-Escalante JC, Comas-Garcia A, Bernal-Silva S, Robles-Espinoza CD, Gomez-Leal G, Noyola DE. 2019. Respiratory syncytial virus A genotype classification based on systematic intergenotypic and intragenotypic sequence analysis. Sci Rep 9:20097.

28. Tramuto F, Maida CM, Mazzucco W, Costantino C, Amodio E, Sferlazza G, Previti A, Immordino P, Vitale F. 2023. Molecular Epidemiology and Genetic Diversity of Human Respiratory Syncytial Virus in Sicily during Pre- and Post-COVID-19 Surveillance Seasons. Pathogens 12.

29. Peret TC, Hall CB, Hammond GW, Piedra PA, Storch GA, Sullender WM, Tsou C, Anderson LJ. 2000. Circulation patterns of group A and B human respiratory syncytial virus genotypes in 5 communities in North America. J Infect Dis 181:1891–6.

30. Ramaekers K, Rector A, Cuypers L, Lemey P, Keyaerts E, Van Ranst M. 2020. Towards a unified classification for human respiratory syncytial virus genotypes. Virus Evol 6:veaa052.

31. Langedijk AC, Harding ER, Konya B, Vrancken B, Lebbink RJ, Evers A, Willemsen J, Lemey P, Bont LJ. 2022. A systematic review on global RSV genetic data: Identification of knowledge gaps. Rev Med Virol 32:e2284.

32. Chen J, Qiu X, Avadhanula V, Shepard SS, Kim DK, Hixson J, Piedra PA, Bahl J. 2022. Novel and extendable genotyping system for human respiratory syncytial virus based on whole-genome sequence analysis. Influenza Other Respir Viruses 16:492–500.

33. Goya S, Galiano M, Nauwelaers I, Trento A, Openshaw PJ, Mistchenko AS, Zambon M, Viegas M. 2020. Toward unified molecular surveillance of RSV: A proposal for genotype definition. Influenza Other Respir Viruses 14:274–285.

34. Nunes D, Vieira C, Sa JM, Araujo GC, Caruso IP, Souza FP. 2022. M2-2 gene as a new alternative molecular marker for phylogenetic, phylodynamic, and evolutionary studies of hRSV. Virus Res 318:198850.

35. Hadfield J, Megill C, Bell SM, Huddleston J, Potter B, Callender C, Sagulenko P, Bedford T, Neher RA. 2018. Nextstrain: real-time tracking of pathogen evolution. Bioinformatics 34:4121–4123.

36. Aksamentov, I, Roemer C., Hodcroft, E. B., Neher, R. A. 2021. Nextclade: clade assignment, mutation calling and quality control for viral genomes. Journal of Open Source Software 6(67), 3773.

37. Gilman MSA, Liu C, Fung A, Behera I, Jordan P, Rigaux P, Ysebaert N, Tcherniuk S, Sourimant J, Eleouet JF, Sutto-Ortiz P, Decroly E, Roymans D, Jin Z, McLellan JS. 2019. Structure of the Respiratory Syncytial Virus Polymerase Complex. Cell 179:193–204 e14.

38. Cao D, Gao Y, Roesler C, Rice S, D’Cunha P, Zhuang L, Slack J, Domke M, Antonova A, Romanelli S, Keating S, Forero G, Juneja P, Liang B. 2020. Cryo-EM structure of the respiratory syncytial virus RNA polymerase. Nat Commun 11:368.

39. Holland LA, Holland SC, Smith MF, Leonard VR, Murugan V, Nordstrom L, Mulrow M, Salgado R, White M, Lim ES. 2023. Genomic Sequencing Surveillance to Identify Respiratory Syncytial Virus Mutations, Arizona, USA. Emerg Infect Dis 29:2380–2382.

40. Goya S, Sereewit J, Pfalmer D, Nguyen TV, Bakhash S, Sobolik EB, Greninger AL. 2023. Genomic Characterization of Respiratory Syncytial Virus during 2022- 23 Outbreak, Washington, USA. Emerg Infect Dis 29:865–868.

41. Rebuffo-Scheer C, Bose M, He J, Khaja S, Ulatowski M, Beck ET, Fan J, Kumar S, Nelson MI, Henrickson KJ. 2011. Whole genome sequencing and evolutionary analysis of human respiratory syncytial virus A and B from Milwaukee, WI 1998–2010. PLoS One 6:e25468.

42. Schobel SA, Stucker KM, Moore ML, Anderson LJ, Larkin EK, Shankar J, Bera J, Puri V, Shilts MH, Rosas-Salazar C, Halpin RA, Fedorova N, Shrivastava S, Stockwell TB, Peebles RS, Hartert TV, Das SR. 2016. Respiratory Syncytial Virus whole-genome sequencing identifies convergent evolution of sequence duplication in the C-terminus of the G gene. Sci Rep 6:26311.

43. Lischer HEL, Shimizu KK. 2017. Reference-guided de novo assembly approach improves genome reconstruction for related species. BMC Bioinformatics 18:474.

44. Bankevich A, Nurk S, Antipov D, Gurevich AA, Dvorkin M, Kulikov AS, Lesin VM, Nikolenko SI, Pham S, Prjibelski AD, Pyshkin AV, Sirotkin AV, Vyahhi N, Tesler G, Alekseyev MA, Pevzner PA. 2012. SPAdes: a new genome assembly algorithm and its applications to single-cell sequencing. J Comput Biol 19:455–77.

45. Sutton TDS, Clooney AG, Ryan FJ, Ross RP, Hill C. 2019. Choice of assembly software has a critical impact on virome characterisation. Microbiome 7:12.

46. Moudy RM, Harmon SB, Sullender WM, Wertz GW. 2003. Variations in transcription termination signals of human respiratory syncytial virus clinical isolates affect gene expression. Virology 313:250–60.

47. Human S, Hotard AL, Rostad CA, Lee S, McCormick L, Larkin EK, Peret TCT, Jorba J, Lanzone J, Gebretsadik T, Williams JV, Bloodworth M, Stier M, Carroll K, Peebles RS, Jr., Anderson LJ, Hartert TV, Moore ML. 2020. A Respiratory Syncytial Virus Attachment Gene Variant Associated with More Severe Disease in Infants Decreases Fusion Protein Expression, Which May Facilitate Immune Evasion. J Virol 95.

48. Kumaria R, Iyer LR, Hibberd ML, Simoes EA, Sugrue RJ. 2011. Whole genome characterization of non-tissue culture adapted HRSV strains in severely infected children. Virol J 8:372.

49. Machkovech HM, Bloom JD, Subramaniam AR. 2019. Comprehensive profiling of translation initiation in influenza virus infected cells. PLoS Pathog 15:e1007518.

50. Girard G, Gultyaev AP, Olsthoorn RC. 2011. Upstream start codon in segment 4 of North American H2 avian influenza A viruses. Infect Genet Evol 11:489–95.

51. Kim HN, Hwang J, Yoon SY, Lim CS, Cho Y, Lee CK, Nam MH. 2023. Molecular characterization of human respiratory syncytial virus in Seoul, South Korea, during 10 consecutive years, 2010-2019. PLoS One 18:e0283873.

52. Cao D, Gao Y, Chen Z, Gooneratne I, Roesler C, Mera C, D’Cunha P, Antonova A, Katta D, Romanelli S, Wang Q, Rice S, Lemons W, Ramanathan A, Liang B. 2023. Structures of the promoter-bound respiratory syncytial virus polymerase. Nature doi:10.1038/s41586-023-06867-y.

53. Liang B, Li Z, Jenni S, Rahmeh AA, Morin BM, Grant T, Grigorieff N, Harrison SC, Whelan SPJ. 2015. Structure of the L Protein of Vesicular Stomatitis Virus from Electron Cryomicroscopy. Cell 162:314–327.

54. Jenni S, Bloyet LM, Diaz-Avalos R, Liang B, Whelan SPJ, Grigorieff N, Harrison SC. 2020. Structure of the Vesicular Stomatitis Virus L Protein in Complex with Its Phosphoprotein Cofactor. Cell Rep 30:53–60 e5.

55. Gould JR, Qiu S, Shang Q, Ogino T, Prevelige PE, Jr., Petit CM, Green TJ. 2020. The Connector Domain of Vesicular Stomatitis Virus Large Protein Interacts with the Viral Phosphoprotein. J Virol 94.

56. Martinello RA, Chen MD, Weibel C, Kahn JS. 2002. Correlation between respiratory syncytial virus genotype and severity of illness. J Infect Dis 186:839–42.

57. Vandini S, Biagi C, Lanari M. 2017. Respiratory Syncytial Virus: The Influence of Serotype and Genotype Variability on Clinical Course of Infection. Int J Mol Sci 18.

58. Felt SA, Achouri E, Faber SR, Lopez CB. 2022. Accumulation of copy-back viral genomes during respiratory syncytial virus infection is preceded by diversification of the copy-back viral genome population followed by selection. Virus Evol 8:veac091.

59. Felt SA, Sun Y, Jozwik A, Paras A, Habibi MS, Nickle D, Anderson L, Achouri E, Feemster KA, Cardenas AM, Turi KN, Chang M, Hartert TV, Sengupta S, Chiu C, Lopez CB. 2021. Detection of respiratory syncytial virus defective genomes in nasal secretions is associated with distinct clinical outcomes. Nat Microbiol 6:672–681.

60. Martin M. 2011. Cutadapt removes adapter sequences from high-throughput sequencing reads. EMBnet Journal 17, n. 1, p. pp. 10-12.

61. Langmead B, Salzberg SL. 2012. Fast gapped-read alignment with Bowtie 2. Nat Methods 9:357–9.

62. Madeira F, Pearce M, Tivey ARN, Basutkar P, Lee J, Edbali O, Madhusoodanan N, Kolesnikov A, Lopez R. 2022. Search and sequence analysis tools services from EMBL-EBI in 2022. Nucleic Acids Res 50:W276–W279.

